# A coarse-grained model for simulations of phosphorylated disordered proteins

**DOI:** 10.1101/2025.03.19.644261

**Authors:** Arriën Symon Rauh, Gustav Stausbøll Hedemark, Giulio Tesei, Kresten Lindorff-Larsen

**Affiliations:** Structural Biology and NMR Laboratory, Linderstrøm-Lang Centre for Protein Science, Department of Biology, University of Copenhagen, Ole Maaløes Vej 5, DK-2200 Copenhagen, Denmark

## Abstract

Protein phosphorylation is a common and essential post-translational modification that affects biochemical properties and regulates biological activities. Phosphorylation is particularly common for intrinsically disordered proteins and can significantly modulate their function and potential to interact with binding partners. To understand the biophysical origins of how phosphorylation of disordered proteins influences their function, it is valuable to investigate how the modifications lead to changes in their conformational ensembles. Here, we have used a top-down data-driven approach to develop a coarse-grained molecular dynamics model compatible with the CALVADOS protein simulation model to study the effects of serine and threonine phosphorylation on the global structural properties of disordered proteins. We parameterise the model using experimental data on the effects of phosphorylation on global dimensions. By comparing with baseline models and simulations using the phosphomimetics aspartate and glutamate, we show that the effect of phosphorylation on the global dimensions of disordered proteins is mostly driven by the additional charge. We envisage that our model can be applied to study the effects of phosphorylation of disordered proteins at the proteome scale as well as to study the important roles of protein phosphorylation on phase separation.

## Introduction

Phosphorylation is the most frequent post-translational modification (PTM) of proteins, where a phosphoryl group is reversibly bound to a side-chain hydroxyl group of a serine (to form pSer), a threonine (pThr), a tyrosine or the imidazole group of a histidine amino acid residue (***Burnett and Kennedy, 1954; Khoury et al., 2011***). This process is driven by protein kinases that catalyse the transfer of a negatively charged phosphoryl group from adenosine-triphosphate to the target amino acid residue and protein phosphatases that facilitate the opposite process. The switch-like reversibility of phosphorylation makes it an ideal mechanism for regulating processes like cell signalling, metabolic pathways and gene transcription (***Wilkinson and Millar, 2000; Humphrey et al., 2015; Viéitez et al., 2022***). Related to this, phosphorylation is also often implicated in diseases where misregulation may cause pathologies (***Cohen, 2001; Šimić et al., 2016; Savastano et al., 2021; Kawahata et al., 2022; Ramasamy et al., 2022; Zhong et al., 2023***).

Phosphorylation is particularly common in and important for modulating the functions of intrinsically disordered proteins and regions (hereafter collectively called IDRs) in processes like transcription and signal processing (***Patwardhan and Miller, 2007; Jin and Gräter, 2021; Newcombe et al., 2022***), and IDRs show enrichment in potential phosphorylation sites (***Iakoucheva, 2004; Ramasamy et al., 2022***). For IDRs, both single and multisite phosphorylation is observed with a variety of functional consequences. Single-site phosphorylation events often result in switch-like functional behaviour where, for example, binding to other proteins (***Solt et al., 2006; Dahal et al., 2017; Sun et al., 2021***) or a membrane is altered (***Harrington et al., 2021***). For multisite phosphorylation, on the other hand, a wider variety of functional effects is known, ranging from switch-like effects with a specific set of phosphorylations to more intricate modulation effects with more complex kinetics (***Salazar and Höfer, 2009***). For example, the consecutive addition of phosphoryl groups at different sites can facilitate an incremental effect, or phosphorylation at different sites can lead to interrelated effects with cooperativity or auto-suppression (***Gingras et al., 1999a***,b; ***Nash et al., 2001; Mittag et al., 2008; Martin et al., 2022***). Additionally, when separate kinases can phosphorylate an IDR, this can result in different phosphorylation patterns that act as barcodes with distinct functional outcomes (***Flotow et al., 1990; Venkatakrishnan et al., 2014; Eiger et al., 2023***). Another interesting specific functional effect of the phosphorylation of IDRs is related to the protein quality control system. When so-called phosphodegrons, a class of short-linear motifs, are phosphorylated, this enables the binding of specific protein ligases, leading to ubiquitylation and subsequent degradation (***Nishizawa et al., 1998; Hunter, 2007; Swaney et al., 2013; Nakagawa and Nakagawa, 2025***).

For IDRs, it can be useful to examine the possible conformational changes caused by the addition of one or more phosphoryl groups to understand how the function changes upon phosphorylation (***Jin and Gräter, 2021; Newcombe et al., 2022; Usher et al., 2024***). For instance, phosphorylation can affect cis/trans isomerisation at proline residues, influence the stability of secondary structural elements, or even induce folding (***Mandell et al., 2007; Bah et al., 2015; Buholzer et al., 2022; Pandey et al., 2023; Bickel and Vranken, 2024; Theisen et al., 2025***). The conformational ensemble of an IDR is encoded in its amino acid sequence composition and patterning (***Van Der Lee et al., 2014; Tesei et al., 2024; Lotthammer et al., 2024***). Phosphorylation causes a change in net charge and charge patterning due to the negative charge of the phosphoryl group at physiological conditions (***Hendus-Altenburger et al., 2019***). When negative charges are added to an IDR with a positive net charge, this can allow for new attractive intra-chain charge-charge interactions which may lead to compaction (***Martin et al., 2016; Gomes et al., 2020; Rieloff and Skepö, 2020; Jin and Gräter, 2021***), whereas an IDR that is already enriched in acidic amino acids may become more extended due to increased repulsive electrostatic interactions (***Gibbs et al., 2017; Kulkarni et al., 2017; Lenton et al., 2017; Jin and Gräter, 2021; Du et al., 2025***). Sequence composition also governs the formation of higher-order assemblies, such as the phase separation (PS) of IDRs into biomolecular condensates (***Pappu et al., 2023***), which may also be modulated by phosphorylation (***Wippich et al., 2013; Monahan et al., 2017; Lu et al., 2018; Guo et al., 2019; Greig et al., 2020***). In the case of homotypic PS where multiple chains of the same IDR phase separate with themselves, the propensity to phase separate generally decreases with increasing absolute net charge per residue (NCPR) of the IDR (***Monahan et al., 2017; Ryan et al., 2021; Jin and Gräter, 2021; Nosella and FormanKay, 2021; Bremer et al., 2022; von Bülow et al., 2024***). Hyperphosphorylation of IDRs like tau or α-synuclein has also been linked to aggregation-related pathologies (***Šimić et al., 2016; Kawahata et al., 2022***).

Experimental techniques such as nuclear magnetic resonance (NMR) spectroscopy, small-angle X-ray scattering (SAXS), and single-molecule Förster resonance energy transfer (smFRET) have been applied to characterise changes in conformational ensembles of IDRs upon phosphorylation (***Martin et al., 2016; Chin et al., 2016; Lenton et al., 2017; Gibbs et al., 2017; Kulkarni et al., 2017; Gomes et al., 2020; Rieloff and Skepö, 2020; Jin and Gräter, 2021; Du et al., 2025; Mittag et al., 2010; Hendus-Altenburger et al., 2017; Mateos et al., 2021***). One complicating factor in the interpretation of these data is that protein kinases used in in-vitro experiments often result in mixtures of IDRs with different phosphorylation states due to the different specificities for the multiple phosphorylation sites (***Gomes et al., 2020; Chen et al., 2025***). To determine the effects of specific phosphorylation patterns, phosphorylation sites can be substituted with aspartate or glutamate residues, which are considered phosphomimetic as their side chains bear a (single) negative charge and have dimensions similar to pSer and pThr (***Dissmeyer and Schnittger, 2011; Hunter, 2012; Chen and Cole, 2015; Monahan et al., 2017***).

Computational methods like Monte Carlo or molecular dynamics (MD) simulations are a useful aid to interpret experiments in integrative studies exploring the conformational dynamics of IDRs (***Thomasen and Lindorff-Larsen, 2022***). Both full-atomistic or coarse-grained (CG) resolutions have been applied in studies of phosphorylation of IDRs (***Rieloff and Skepö, 2020; Perdikari et al., 2021; Jin and Gräter, 2021; Ranganathan et al., 2023; Changiarath et al., 2024; Usher et al., 2024; Zippo et al., 2024***). For example, ***Jin and Gräter*** (***2021***) used atomistic MD simulations to study the effects of multisite phosphorylation on Ash1, the C-terminal domain of RNA polymerase II subunit RPB1 (CTD2), the E-Cadherin intracellular domain, and a p130Cas fragment. Alternatively, Monte Carlo simulations using the implicit-solvent all-atom ABSINTH force field have been used to explore the changes in intra-chain interactions caused by phosphomimetics (***Martin et al., 2016***) or phosphorylation (***Usher et al., 2024***). Residue-level CG MD approaches have also been applied to efficiently study global dimensions and PS of phosphorylated IDRs (***Dignon et al., 2018a; Dannenhoffer-Lafage and Best, 2021; Regy et al., 2021; Tesei et al., 2021; Joseph et al., 2021; Latham and Zhang, 2021; Jung et al., 2024; Jussupow et al., 2024; Lohberger et al., 2025***). ***Perdikari et al. (2021)*** introduced pSer and pThr into the hydrophobicity scale (HPS) model (***Dignon et al., 2018a)***, which has been applied to study the effects of phosphorylation on the PS of IDRs like FUS LCD, LAF-1, and CTD2 (***Perdikari et al., 2021; Ranganathan et al., 2023; Changiarath et al., 2024)***. Further, ***Zippo et al. (2024)*** recently implemented this model in a non-equilibrium simulation approach that can be used to explore the phosphorylation kinetics in a biomolecular condensate.

We previously presented the residue-level CG model CALVADOS to predict conformational and phase properties of IDRs from amino acid sequence and solution conditions (***Tesei et al., 2021; Tesei and Lindorff-Larsen, 2023)***. The model builds upon previous HPS-type models where residues are assigned a λ value describing their amino acid-specific hydropathy (***Dignon et al., 2018b; Regy et al., 2021)***, and we reparameterised the hydrophobicity (‘stickiness’) scale against single-chain data from SAXS and NMR experiments using a Bayesian parameter-learning approach (***Norgaard et al., 2008)***. In this model, all amino acid residues are represented by a single bead and residue– residue nonbonded interactions are determined by a combination of the stickiness parameters and salt-screened electrostatics.

We have used our model to study the conformational dynamics of the human intrinsically disordered proteome (***Tesei et al., 2024)***, to train predictive models for global dimensions and phase separation propensities (***Tesei et al., 2024; von Bülow et al., 2024)***, and to design disordered proteins with targeted properties (***Pesce et al., 2024)***. In addition to IDRs, CALVADOS has been expanded to model multi-domain proteins (***Cao et al., 2024)***, RNA (***Yasuda et al., 2025)***, and crowding effects with poly-ethylene glycol (***Rauh et al., 2025)***. Recently, the Perdikari parameters (***Perdikari et al., 2021)*** have been used with CALVADOS to study the effect of phosphorylation of Tau (***Lohberger et al., 2025)***.

To enable the study of phosphorylation effects on the global conformational dynamics and PS behaviour of IDRs with our CALVADOS model, we here implemented a model for phosphorylated serine (pSer) and threonine (pThr) based on single-chain compaction data. The model is thus aimed at capturing changes in the global structural properties and not, for example, changes in local helix formation. In line with the general strategy of parameterising CALVADOS, we target primarily experimental data on proteins. We therefore first collect a dataset of SAXS measurements of the dimensions of IDRs both in their unphosphorylated and phosphorylated states (***Martin et al., 2016; Chin et al., 2016; Lenton et al., 2017; Gibbs et al., 2017; Kulkarni et al., 2017; Gomes et al., 2020; Rieloff and Skepö, 2020; Jin and Gräter, 2021; Du et al., 2025)***. We then use this dataset to parameterise our model, evaluating the change in compaction upon phosphorylation. We also use the model to compare conformational ensembles of phosphorylated proteins with those with phosphomimetic residues. In line with previous findings (***Jin and Gräter, 2021)***, we observe that the largest impact of phosphorylation on the conformational ensembles can be attributed to the introduction of additional negative charges, whereas changes in hydropathy lead to subtler effects.

## Results and Discussion

### Dataset of phosphorylated intrinsically disordered regions

In this work, we aim to introduce phosphorylated serine (pSer) and threonine (pThr) residues into the CALVADOS protein model, using experimental data to inform their parameterisation. To this end, we set out to curate a training set of proteins for which chain compaction data is available for both phosphorylated and unphosphorylated states (Figure 1A and B). We explored the literature and identified nine different IDRs with in total 11 different phosphorylation states for which either SAXS or smFRET data have been reported (Figure 1A, and Tables S1, S2, S3, S4, S5, and Table S6): A C-terminal IDR from the *S. cerevisiae* transcription factor Ash1 (Ash1) (***Martin et al., 2016; Jin and Gräter, 2021)***; the C-terminal IDR of RNA polymerase II subunit RPB1 (CTD2) (***Gibbs et al., 2017; Jin and Gräter, 2021)***; the N-terminal IDR of the Sic1 protein (Sic1) (***Gomes et al., 2020)***; the IDR Prostate-Associated Gene 4 (PAGE4) (***Kulkarni et al., 2017)***; 15-residue-long N-terminal fragment of the IDR Statherin (SN15) (***Rieloff and Skepö, 2020)***; a recombinant Osteopontin peptide (rOPN) (***Lenton et al., 2017)***; the N-terminal disordered transactivation domain of the oestrogen receptor (ERα) (***Du et al., 2025)***; and two versions of a short Tau peptide with either serines (TauS) or threonines (TauT) (***Chin et al., 2016)***. Part of this dataset overlaps with the test proteins used by ***Usher et al. (2024)*** to implement the OPLS parameters for pSer and pThr in the ABSINTH model.

**Figure 1.**
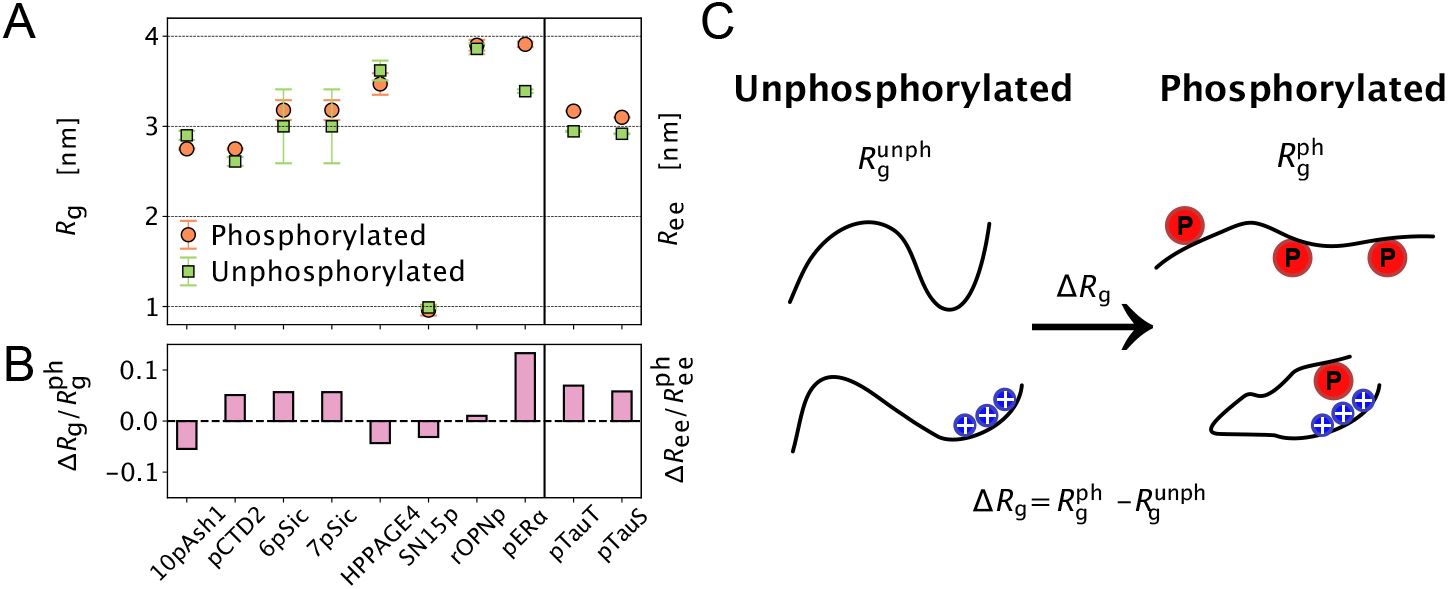
Dataset of global dimensions of phosphorylated intrinsically disordered regions. **(A)** Experimental radii of gyration, *R*_*g*_ (*left*), derived from SAXS and end-to-end distances, *R*_ee_ (*right*), derived from smFRET of phosphorylated (orange circles) and unphosphorylated (green squares) IDRs. (**B**) Relative change in global dimensions upon phosphorylation, 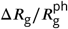 (*left*) and 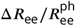 (*right*), with respect to the phosphorylated IDR. (**C**) Graphical illustration of phosphorylation-induced conformational changes. *Top*, phosphorylation along the sequence causes chain expansion due to repulsive interactions between the negatively charged phosphoryl groups. *Bottom*, phosphorylation promotes chain compaction due to attractive charge-charge interactions.

For both Sic1 and PAGE4, we have two different phosphorylated constructs. Based on mass spectrometry, ***Gomes et al. (2020)*** reported a mixture of three different populations of phosphorylated Sic1. Of these, the two major populations are a 6-fold phosphorylated (6pSic1) and a 7-fold fully phosphorylated (7pSic1) Sic1. We decided to include both 6pSic1 and 7pSic1 as separate entries in our dataset since their contributions to the ensemble-averaged experimental *R*_g_ are considered to have similar weights. For PAGE4, ***Kulkarni et al. (2017)*** conducted experiments using two different kinases, which resulted in two different phosphorylation patterns with functional consequences for the binding of PAGE4 to transcription factor AP-1. Phosphorylation of PAGE4 by homeodomain-interacting protein kinase 1 (HIPK1) occurs at two positions and results in HPPAGE4, which has a more compact conformational ensemble than unphosphorylated PAGE4. Conversely, CDC-like kinase 2 (CLK2) phosphorylates PAGE4 at six additional sites and the resulting CPPAGE4 is considerably more expanded (Tables S3, S4, and S6). We decided not to include CPPAGE4 in our dataset because of the high reported error on the experimental *R*_g_. Further, for 10pAsh1 and pCTD2, we chose to use the *R*_g_ values obtained by ***Jin and Gräter*** (***2021)*** from their analysis of the original SAXS data.

### Parameterisation of phosphorylated beads

After compiling the dataset of phosphorylated IDRs, we set out to fine-tune the CALVADOS model parameters by comparison with experiments. First, we estimate the bead sizes of pSer and pThr by increasing the volume of the unphosphorylated residues by 0.041 nm^3^ (***Newcombe et al., 2022)***. Second, we calculate the charge of pSer and pThr based on solution pH and using the experimental p*K*_*a*_ ≈ 6 (Eq. 7) for the secondary ionization of pSer and pThr in random coils (***Hendus-Altenburger et al., 2019)***.

We then parameterised the stickiness of pSer (λ_pSer_) and pThr (λ_pThr_). Given the limited amount of available experimental data, we chose to optimise a single parameter (Δλ) that reports on the change in stickiness upon phosphorylation. This approach implies that the addition of a phosphoryl group causes the same shift in λ for both serine and threonine residues. We therefore scanned Δλ and evaluated the agreement between experiments and single-chain simulations performed with different stickiness values, λ_pX_ = λ_X_ + Δλ. To quantify this agreement, we use the ratio of the change in *R*_g_ to the *R*_g_ of the phosphorylated protein, 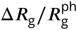. Our parameter scan shows that the agreement with experimental data increases with decreasing Δλ (Figures 2A, S1, S2 and Table S6). As the final value, we choose Δλ = −0.37, as it corresponds to setting λ_pThr_ ≈ 0. Lower values would result in negative stickiness, which would make the Ashbaugh-Hatch potential physically inconsistent (***Ashbaugh and Wood, 1997; Ashbaugh and Hatch, 2008)***.

**Figure 2.**
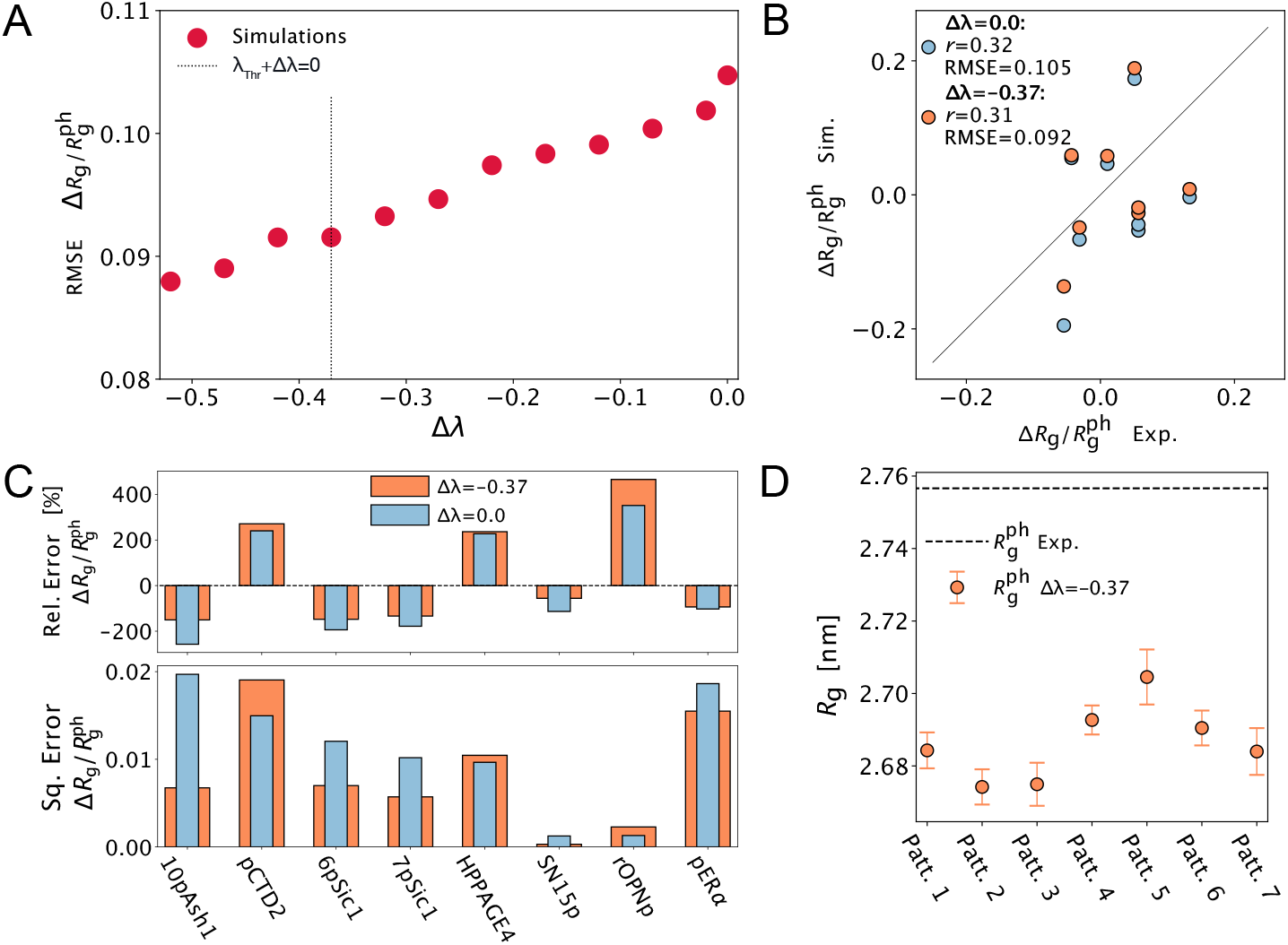
Scanning a Δλ parameter to reproduce changes in global dimensions of IDRs upon phosphorylation. (**A**) Root-mean-square error (RMSE) with respect to experiments of the predicted relative changes in *R*_*g*_ upon phosphorylation, 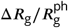, as a function of Δλ. The vertical dotted line shows the Δλ resulting in λ_pThr_ ≈ 0. (**B**) Comparison between simulated and experimental 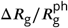 for Δλ = 0.0 (blue) and Δλ = −0.37 (orange). The solid black line shows the diagonal and the legend reports Pearson correlation coefficients, *r*, and RMSEs. (**C**) Relative errors (*top*) and squared errors (*bottom*) per protein shown for Δλ = 0.0 (narrow blue bar) and Δλ = −0.37 (wide orange bar). (**D**) Comparison of the simulated 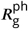 for 6pSic1 with all possible phosphorylation patterns simulated with Δλ = −0.37 (circles). The dashed line shows the experimental 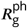

To establish the effect of lowering λ_pThr_ and λ_pSer_, in addition to the effective change in stickiness caused by increasing the size of the beads, we examined the differences between results with Δλ = 0.0 and Δλ = −0.37. The comparison indicates that, overall, the trends remain consistent between the two models and that the improved agreement with experimental data for Δλ = −0.37 (Figures 2B–C and S3) is primarily due to 10pAsh1, with smaller contributions from pSic1 and pERα. The pronounced sensitivity to the change in hydropathy of 10pAsh1 can be explained by its hyperphosphorylation (12.3 % of its residues are phosphorylated, see Table S3) and its compact conformational ensemble (Table S7).

We also tested to what extent the phosphorylation pattern along the linear sequence influences compaction, thus probing the importance of changes in charge patterning upon phosphorylation. This is specifically important for phosphorylated Sic1, for which ***Gomes et al. (2020)*** report two main populations with 6 and 7 phosphorylated residues. The presence of an unphosphorylated phosphosite in 6-fold pSic1 (6pSic1) makes the exact phosphorylation pattern uncertain. To explore potential differences between different positions of the 6 phosphorylated serines, we performed single-chain simulations for all 7 phosphorylation variants of 6pSic1, using Δλ = −0.37. The simulations show that the differences in compaction between the phosphorylation variants of 6pSic1 are small with the *R*_g_ values for these variants covering a range of 0.03 nm with a mean value of 2.69 nm (Figure 2D).

### Comparison of the phosphorylation model to alternative models

Having established a model for both pSer and pThr, we compared it to two alternative models. First, we compared our data-driven model (Δλ = −0.37) with a model where we simply add equivalent negative charges as pSer and pThr to serine (Ser^≈−2^) and threonine (Thr^≈−2^) beads, without modifying their size or stickiness (“charge model”). This comparison captures the effect of the modified charges relative to the other modified parameters. We therefore performed single-chain simulations of all the proteins with Ser^≈−2^ and Thr^≈−2^ instead of pSer and pThr. The high correlation between changes in compaction predicted by the charge model and by the data-driven model shows that a substantial part of the performance of our model can be explained by the modified charges (Figures 3A and S4A). We observe larger differences in performance for IDRs that become more compact upon phosphorylation, and especially for 10pAsh1, which is the most sensitive to the hydropathy correction.

**Figure 3.**
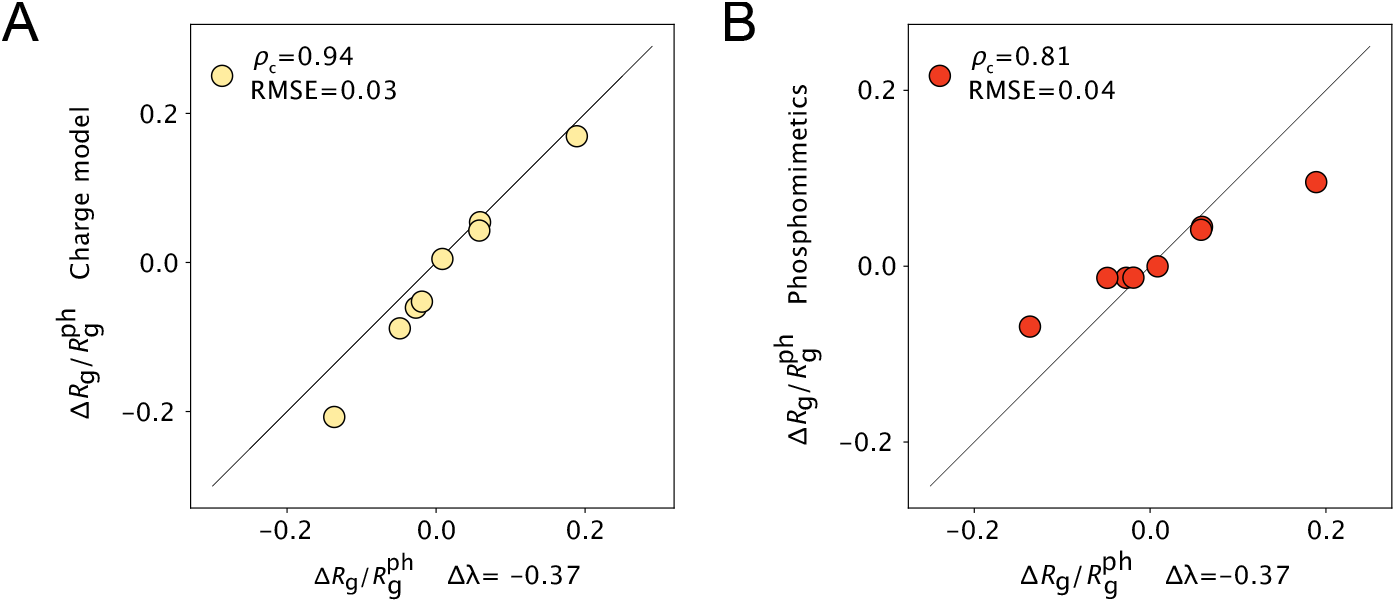
Comparison of the phosphorylation model with alternative models. (**A**) Comparison of 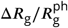 from simulations with Δλ = −0.37 and from simulations where serine and threonine residues at the phosphorylation sites are replaced with charged versions of Ser and Thr, respectively, which have a charge of ≈ −2 and the same bead sizes and stickiness as the unphosphorylated residues. (**B**) Comparison of 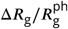 from simulations with Δλ = −0.37 and from simulations where serine and threonine residues at the phosphorylation sites are replaced with the commonly used phosphomimetics aspartate and glutamate residues, respectively. Black solid lines show the diagonals, and we report the concordance correlation coefficient, *ρ*_c_, (***Lin, 1989)*** and the RMSE in the legends.

Second, we compared simulations of our data-driven model for phosphorylated IDRs with phosphomimetic variants, where pSer and pThr are modelled as Asp and Glu, respectively. This enables us better to capture the effect of the charge valency, while accounting for differences in size and stickiness because in the CALVADOS model Glu is larger than Asp and both have low stickiness. This analysis shows that phosphomimetic variants yield smaller changes in compaction compared to phosphorylation, as captured by our data-driven model. (Figures 3B and S4B). This trend is expected since single negative charges introduced in the phosphomimetic variants result in weaker charge-charge interactions than those of the divalent charges of the phosphorylated residues.

## Conclusion

In this work, we set out to implement a model for phosphorylated IDRs which is compatible with the CALVADOS 2 CG model (***Tesei and Lindorff-Larsen, 2023)***. To do so, we first compiled a dataset of IDRs with experimental data reporting on their global dimensions in both phosphorylated and unphosphorylated states (***Chin et al., 2016; Martin et al., 2016; Gibbs et al., 2017; Kulkarni et al., 2017; Lenton et al., 2017; Gomes et al., 2020; Rieloff and Skepö, 2020; Jin and Gräter, 2021; Du et al., 2025)***. We then used this dataset to determine the change in stickiness of Ser and Thr residues and found that Δλ = −0.37—corresponding to a stickiness value of 0 for pThr and of 0.09 for pSer—leads to improved agreement with experiments. We note that this model only captures effects on global structural properties (as measured by SAXS and smFRET experiments) and not, for example, changes to local structure (***Newcombe et al., 2022)***. To shed further insight into what drives changes in global compaction, we compared the model with two alternative approaches: (i) a naive model where we add charges to the standard CALVADOS Ser and Thr beads and (ii) phosphomimetic variants. These analyses show that the effect of phosphorylation on compaction can be mainly attributed to the increased negative charge of the phosphoryl group, whereas the change in hydropathy contributes to a lesser extent. For Ash1 we find that the change in stickiness is necessary to better capture the experimental change in *R*_g_. We also note that phosphomimetic variants (with charge −1) tend to underestimate the effect of phosphorylation (with charge around −2) on IDR dimensions, in agreement with the observed predominant role of the charge of the phosphoryl group (***Dissmeyer and Schnittger, 2011; Hunter, 2012; Chen and Cole, 2015)***.

In contrast to previously developed residue-level (***Perdikari et al., 2021)*** and implicit solvent allatom models (***Usher et al., 2024)***, we here employed a top-down approach to fine-tune our model parameters. Due to the limited amount of experimental data, we chose to optimise a single parameter representing the change in stickiness of Ser and Thr upon the addition of a phosphate group. In comparison to a two-parameter fit, this simplification may limit the ability of the model to capture subtler differences between pSer and pThr (***Newcombe et al., 2022)***. To overcome this limitation and potentially derive parameters also for phospho-tyrosine, future work could include measurements from paramagnetic relaxation enhancement NMR spectroscopy (***Mittag et al., 2010; Hendus-Altenburger et al., 2017; Mateos et al., 2021)***. We note that the bottom-up and top-down approaches lead to comparable parameters (Table S8), suggesting that they may be combined.

In addition to the study of phosphorylation effects on single-chain ensemble properties, our model will be useful to investigate the influence of phosphorylation on the phase separation of IDRs (***Wippich et al., 2013; Monahan et al., 2017; Lu et al., 2018; Guo et al., 2019; Greig et al., 2020; Changiarath et al., 2024)***. Moreover, one may use our amino acid-specific parameters as input for machine learning-based predictors for compaction (***Tesei et al., 2024)*** and PS propensity (***von Bülow et al., 2024)*** since these models use physics-based features that include charge and amino-acid stickiness. Future work could include studying a large variety of proteins with different phosphorylation patterns. Finally, our model can be used in a non-equilibrium simulation setup to explore how proteins in condensates are phosphorylated (***Zippo et al., 2024)***, for example, probing for processivity (***Patwardhan and Miller, 2007)***. Thus, we envisage that the ability to include effects of serine and threonine phosphorylation in simulations of IDRs will enable a broader set of studies of the biophysical and biological consequences of protein phosphorylation.

## Methods and Materials

### CALVADOS simulations

We performed all molecular dynamics simulations using openMM (v7.5) (***Eastman et al., 2017)*** in the NVT ensemble using a Langevin integrator with a time step of 10 fs and a friction coefficient of 0.01 ps^−1^. As force field, we used the residue level CALVADOS 2 model (***Tesei and Lindorff-Larsen, 2023)***. We model bonds with a harmonic potential,

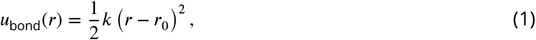

where *k* = 8030 kJ mol^−1^ nm^−2^, *r*_0_ = 0.38 nm. We model non-ionic interactions with a truncated and shifted Ashbaugh-Hatch potential (***Ashbaugh and Wood, 1997; Ashbaugh and Hatch, 2008)***,

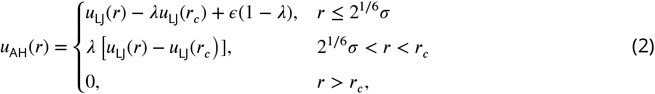

where *ϵ* = 0.8368 kJ mol^−1^, *r*_*c*_ = 2 nm, and λ is the arithmetic average of the stickiness values of the two interacting residues. *u*_LJ_ is the 12-6 Lennard-Jones potential,

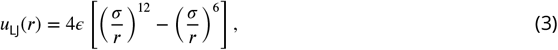

where *σ* is the arithmetic average of the Van der Waals diameters of the interacting residues with amino acid-specific values determined by ***Kim and Hummer*** (***2008)***. We model salt-screened ionic interactions with a Debye-Hückel potential truncated and shifted at a cut-off of *r*_*c*_ = 4 nm,

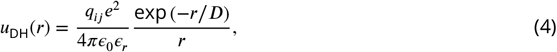

where *q*_*ij*_ is the product of the charge numbers of the interacting residues, *e* is the elementary charge, *ϵ*_0_ is the vacuum permittivity, and 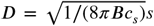 is the Debye length of an electrolyte solution of ionic strength, *c*_*s*_, and Bjerrum length, *B*. We model the temperature-dependent dielectric constant of the implicit aqueous solution, *ϵ*_*r*_, with an empirical relationship (***Akerlof and Oshry, 1950)***,

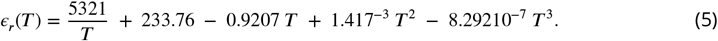

We use the Henderson-Hasselbalch equation (***Henderson, 1908; Hasselbalch, 1916; Po and Senozan, 2001)*** to calculate the pH-dependent partial charges on histidine as

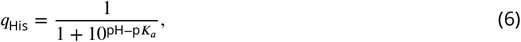

with p*K*_*a*_ = 6.00, and the partial charges on phosphorylated serine and threonine residues as

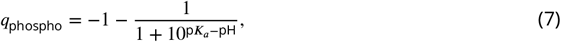

with p*K*_*a*_ = 6.01 for pSer and p*K*_*a*_ = 6.30 for pThr (***Hendus-Altenburger et al., 2019)***.

### Parameterisation of the phosphorylation model

To parameterise the stickiness of phospho-serine (λ_pSer_) and phospho-threonine (λ_pThr_) beads in the CALVADOS model, we determine a value for Δλ in a parameter scan that spans values from 0.0 to −0.52, representing the change in hydropathy caused by the addition of the phosphoryl group. For every value of Δλ, we run single-chain simulations of the phosphorylated IDRs in a cubic box with the simulation settings outlined in Table 1 (see Table S1 for the sequences of both phosphorylated and unphosphorylated IDRs). Additionally, we run single-chain simulations of the unphosphorylated IDRs with the same simulation settings. We run three replicates of every system to collect approximately 5000 independent frames.

**Table 1.** Simulation-box side length, *L*_*xyz*_, temperature, *T*, ionic strength, *c*_*s*_, and pH for phosphorylated and unphosphorylated IDRs simulated in this study.

| Phosphorylated protein | Unphosphorylated protein | $L_{xyz}$ [nm] | $T$ [K] | $c_s$ [mM] | pH | Source |
| --- | --- | --- | --- | --- | --- | --- |
| 10pAsh1 | Ash1 | 34.40 | 298 | 190 | 7.5 | (Martin et al., 2016) |
| pCTD2 | CTD2 | 35.16 | 296 | 117 | 7.5 | (Gibbs et al., 2017) |
| 6pSic1 | Sic1 | 38.58 | 293 | 191 | 7.5 | (Gomes et al., 2020) |
| 7pSic1 | Sic1 | 38.58 | 293 | 191 | 7.5 | (Gomes et al., 2020) |
| HPPAGE4 | WTPAGE4 | 42.38 | 293 | 125 | 7.8 | (Kulkarni et al., 2017) |
| CPPAGE4 | WTPAGE4 | 42.38 | 293 | 125 | 7.8 | (Kulkarni et al., 2017) |
| SN15p | SN15n | 15.00 | 293 | 166 | 7.5 | (Rieloff and Skepö, 2020) |
| rOPNp | rOPN | 65.56 | 293 | 430 | 7.5 | (Lenton et al., 2017) |
| pERα | ERα | 73.54 | 278 | 200 | 7.4 | (Du et al., 2025) |
| pTauT | TauT | 15.00 | 293 | 0.095 | 6.8 | (Chin et al., 2016) |
| pTauS | TauS | 15.00 | 293 | 0.095 | 6.8 | (Chin et al., 2016) |

To evaluate which Δλ value best reproduces the experimental data, we analysed the relative change in compaction upon phosphorylation, 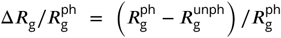. To quantify the agreement with experiments, we calculate the relative error,

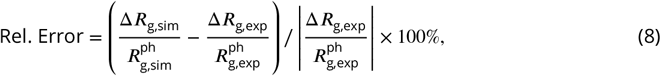

the squared error,

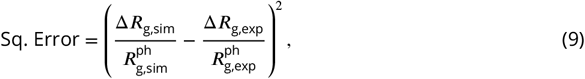

and the root-mean-squared error with respect to the experimental data of the relative change in compaction for the *n* proteins in our dataset,

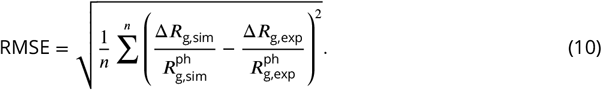

## Data and Code Availabillity

All the data and code used for this work are available via https://github.com/KULL-Centre/_2025_rauh_phosphorylation. The phosphorylation model is available in the CALVADOS package https://github.com/KULL-Centre/CALVADOS.

## Acknowledgements

We thank the members of SBiNLab for helpful discussions. We acknowledge access to computational resources from the ROBUST Resource for Biomolecular Simulations (supported by the Novo Nordisk Foundation grant no. NF18OC0032608). This work is a contribution from the PRISM (Protein Interactions and Stability in Medicine and Genomics) centre funded by the Novo Nordisk Foundation (to K.L.-L.; NNF18OC0033950).

## Competing Interests

K.L.-L. holds stock options in and is a consultant for Peptone. The remaining authors declare no competing interests.

## Supplemental Information

**Table S1.**
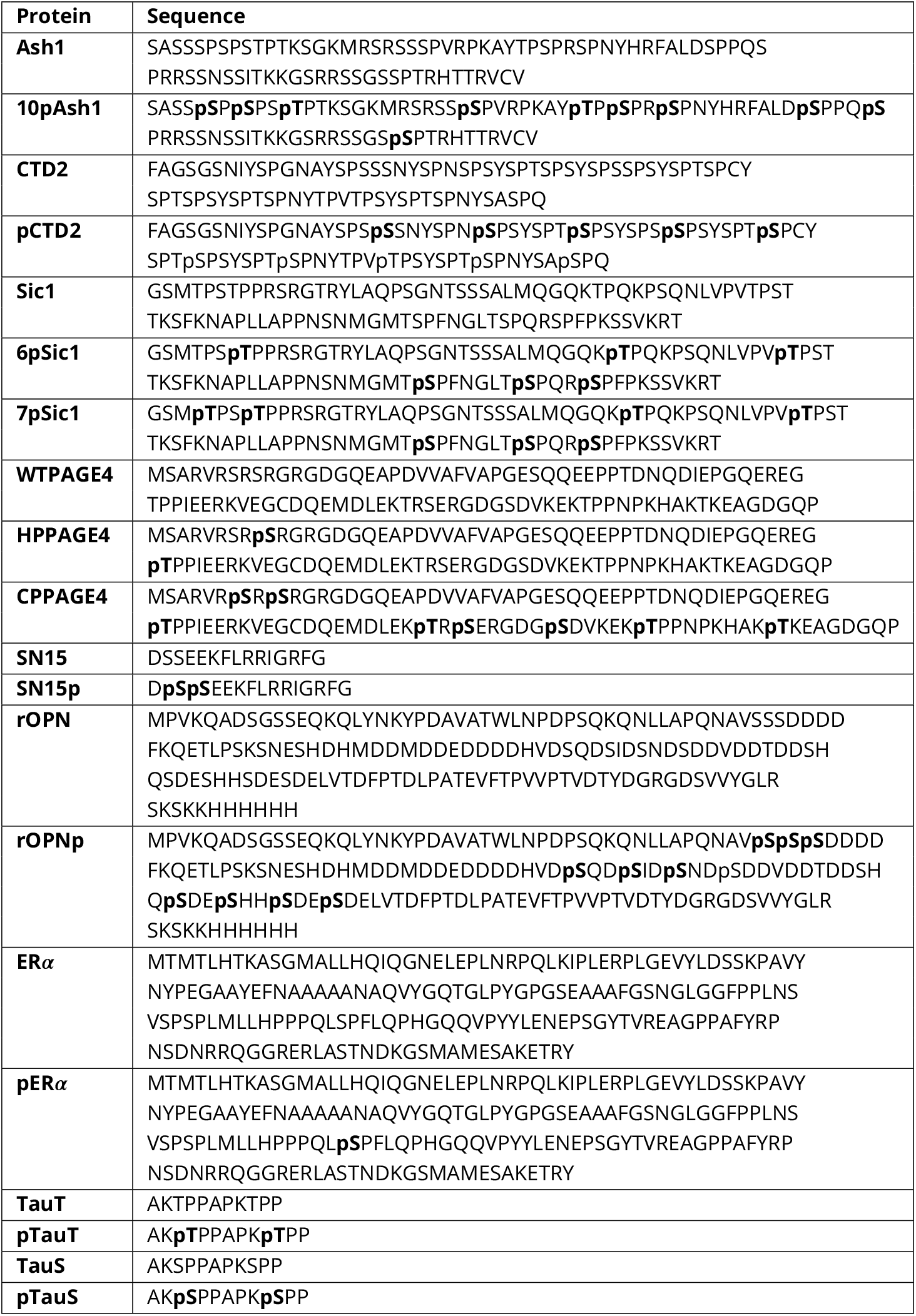
Sequences of unphosphorylated and phosphorylated proteins simulated in this study. In every sequence, the phosphorylation sites are emphasised with bold text.

**Table S2.**
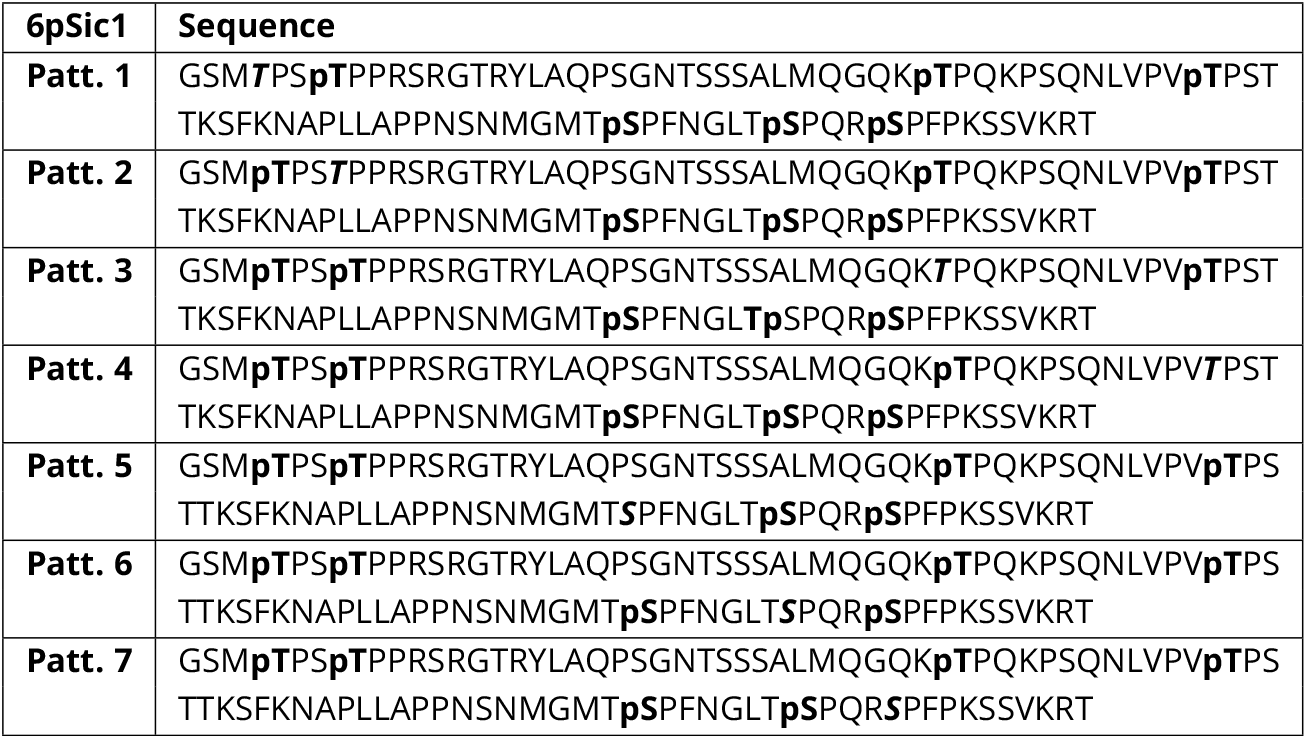
Sequences of the different phosphorylation patterns for the 6-fold phosphorylated Sic1 (6pSic1). In every sequence, the phosphorylation sites are emphasised with bold text and the unphosphorylated site is emphasised with bold and italic text.

**Table S3.** Fractions of phosphorylated serine (*f*_pSer_) and threonine (*f*_pThr_) residues, *f*_ph_ = *f*_pSer_ + *f*_pThr_, and sequence lengths, *N*, of the proteins simulated in this study.

| Protein | $f_{pSer}$ | $f_{pThr}$ | $f_{ph}$ | $N$ |
| --- | --- | --- | --- | --- |
| 10pAsh1 | 0.10 | 0.02 | 0.12 | 81 |
| pCTD2 | 0.11 | 0.01 | 0.12 | 83 |
| 6pSic1 | 0.03 | 0.03 | 0.07 | 92 |
| 7pSic1 | 0.03 | 0.04 | 0.08 | 92 |
| HPPAGE4 | 0.01 | 0.01 | 0.02 | 102 |
| CPPAGE4 | 0.04 | 0.04 | 0.08 | 102 |
| SN15p | 0.13 | 0.00 | 0.13 | 15 |
| rOPNp | 0.07 | 0.00 | 0.07 | 163 |
| pERα | 0.01 | 0.00 | 0.01 | 184 |
| pTauT | 0.00 | 0.18 | 0.18 | 11 |
| pTauS | 0.18 | 0.00 | 0.18 | 11 |

**Table S4.** Radii of gyration, *R*_g_, for phosphorylated and unphosphorylated IDRs measured by SAXS experiments.

| Protein<br>phos. | $R_{g,exp}^{ph}$<br>[nm] | $\sigma_{R_{g,exp}^{ph}}$<br>[nm] | Protein<br>unph. | $R_{g,exp.}^{unph}$<br>[nm] | $\sigma_{R_{g,exp}^{unph}}$<br>[nm] | source |
| --- | --- | --- | --- | --- | --- | --- |
| 10pAsh1 | 2.75 | 0.01 | Ash1 | 2.9 | 0.05 | (Martin et al., 2016) |
| pCTD2 | 2.75 | 0.01 | CTD2 | 2.61 | 0.05 | (Gibbs et al., 2017) |
| 6pSic | 3.18 | 0.11 | Sic1 | 3.00 | 0.41 | (Gomes et al., 2020) |
| 7pSic | 3.18 | 0.11 | Sic1 | 3.00 | 0.41 | (Gomes et al., 2020) |
| HPPAGE4 | 3.47 | 0.12 | HPPAGE4 | 3.62 | 0.11 | (Kulkarni et al., 2017) |
| CPPAGE4 | 4.98 | 0.19 | CPPAGE4 | 3.62 | 0.11 | (Kulkarni et al., 2017) |
| SN15p | 0.96 | 0.06 | SN15n | 0.99 | 0.01 | (Rieloff and Skepö, 2020) |
| rOPNp | 3.9 | 0.06 | rOPN | 3.86 | 0.06 | (Lenton et al., 2017) |
| pERα | 3.91 | 0.03 | ERα | 3.39 | 0.02 | (Du et al., 2025) |

**Table S5.** End-to-end distances, *R*_ee_, of phosphorylated and unphosphorylated Tau peptides with either pThr and pSer measured by smFRET experiments (***Chin et al., 2016)***.

| Protein<br>phos. | $R_{ee,exp}^{ph}$<br>[nm] | $\sigma_{R_{ee,exp}^{ph}}$<br>[nm] | Protein<br>unph. | $R_{ee,exp}^{unph}$<br>[nm] | $\sigma_{R_{ee,exp}^{unph}}$<br>[nm] |
| --- | --- | --- | --- | --- | --- |
| pTauT | 3.169 | 0.004 | TauT | 2.945 | 0.007 |
| pTauS | 3.100 | 0.004 | TauS | 2.917 | 0.005 |

**Table S6.**
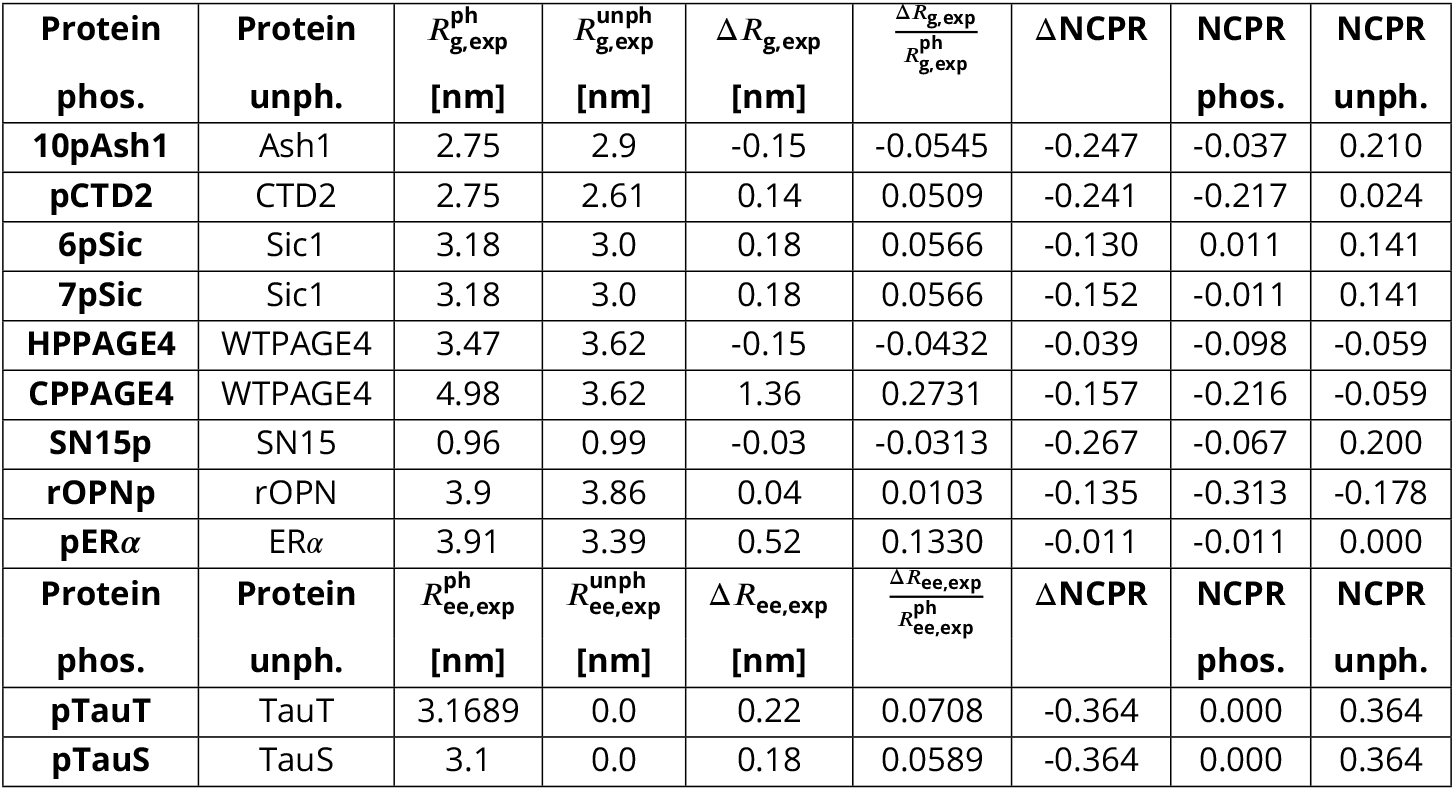
Overview of experimental data reporting on the global chain dimensions and the net charge per residue (NCPR).

| **Protein**<br>**phos.** | **Protein**<br>**unph.** | $R_{g,exp}^{ph}$<br>[nm] | $R_{g,exp}^{unph}$<br>[nm] | $\Delta R_{g,exp}$<br>[nm] | $\frac{\Delta R_{g,exp}}{R_{g,exp}^{ph}}$ | $\Delta NCPR$ | **NCPR**<br>**phos.** | **NCPR**<br>**unph.** |
| --- | --- | --- | --- | --- | --- | --- | --- | --- |
| **10pAsh1** | Ash1 | 2.75 | 2.9 | -0.15 | -0.0545 | -0.247 | -0.037 | 0.210 |
| **pCTD2** | CTD2 | 2.75 | 2.61 | 0.14 | 0.0509 | -0.241 | -0.217 | 0.024 |
| **6pSic** | Sic1 | 3.18 | 3.0 | 0.18 | 0.0566 | -0.130 | 0.011 | 0.141 |
| **7pSic** | Sic1 | 3.18 | 3.0 | 0.18 | 0.0566 | -0.152 | -0.011 | 0.141 |
| **HPPAGE4** | WTPAGE4 | 3.47 | 3.62 | -0.15 | -0.0432 | -0.039 | -0.098 | -0.059 |
| **CPPAGE4** | WTPAGE4 | 4.98 | 3.62 | 1.36 | 0.2731 | -0.157 | -0.216 | -0.059 |
| **SN15p** | SN15 | 0.96 | 0.99 | -0.03 | -0.0313 | -0.267 | -0.067 | 0.200 |
| **rOPNp** | rOPN | 3.9 | 3.86 | 0.04 | 0.0103 | -0.135 | -0.313 | -0.178 |
| **pERα** | ERα | 3.91 | 3.39 | 0.52 | 0.1330 | -0.011 | -0.011 | 0.000 |
| **Protein**<br>**phos.** | **Protein**<br>**unph.** | $R_{ee,exp}^{ph}$<br>[nm] | $R_{ee,exp}^{unph}$<br>[nm] | $\Delta R_{ee,exp}$<br>[nm] | $\frac{\Delta R_{ee,exp}}{R_{ee,exp}^{ph}}$ | $\Delta NCPR$ | **NCPR**<br>**phos.** | **NCPR**<br>**unph.** |
| **pTauT** | TauT | 3.1689 | 0.0 | 0.22 | 0.0708 | -0.364 | 0.000 | 0.364 |
| **pTauS** | TauS | 3.1 | 0.0 | 0.18 | 0.0589 | -0.364 | 0.000 | 0.364 |

**Table S7.** Apparent Flory scaling exponents estimated from simulations as detailed by ***Tesei et al. (2024)***. Values are reported as mean ± standard deviation over three independent simulation replicas.

| Protein | $\nu_{\text{unph}}$ | $\nu_{\text{ph}}^{\Delta\lambda=-0.37}$ | $\nu_{\text{ph}}^{\Delta\lambda=0.0}$ | $\nu_{\text{ch.mod.}}^{\text{ph}}$ | $\nu_{\text{ph.mim.}}^{\text{ph}}$ |
| --- | --- | --- | --- | --- | --- |
| **10pAsh1** | $0.567 \pm 0.001$ | $0.501 \pm 0.001$ | $0.478 \pm 0.002$ | $0.484 \pm 0.001$ | $0.535 \pm 0.001$ |
| **pCTD2** | $0.514 \pm 0.001$ | $0.580 \pm 0.003$ | $0.574 \pm 0.003$ | $0.576 \pm 0.003$ | $0.546 \pm 0.002$ |
| **6pSic1** | $0.543 \pm 0.001$ | $0.525 \pm 0.001$ | $0.514 \pm 0.001$ | $0.511 \pm 0.001$ | $0.535 \pm 0.001$ |
| **7pSic1** | $0.543 \pm 0.001$ | $0.527 \pm 0.001$ | $0.517 \pm 0.001$ | $0.516 \pm 0.002$ | $0.532 \pm 0.001$ |
| **HPPAGE4** | $0.510 \pm 0.003$ | $0.541 \pm 0.003$ | $0.538 \pm 0.003$ | $0.536 \pm 0.003$ | $0.535 \pm 0.003$ |
| **CPPAGE4** | $0.510 \pm 0.003$ | $0.572 \pm 0.003$ | $0.570 \pm 0.003$ | $0.572 \pm 0.003$ | $0.557 \pm 0.003$ |
| **SN15p** | $0.574 \pm 0.008$ | $0.521 \pm 0.003$ | $0.501 \pm 0.003$ | $0.491 \pm 0.004$ | $0.561 \pm 0.005$ |
| **rOPNp** | $0.544 \pm 0.002$ | $0.540 \pm 0.003$ | $0.538 \pm 0.003$ | $0.533 \pm 0.003$ | $0.541 \pm 0.003$ |
| **pERα** | $0.484 \pm 0.001$ | $0.487 \pm 0.001$ | $0.483 \pm 0.001$ | $0.489 \pm 0.001$ | $0.482 \pm 0.001$ |
| **pTauT** | $0.611 \pm 0.006$ | $0.624 \pm 0.006$ | $0.613 \pm 0.006$ | $0.600 \pm 0.007$ | $0.617 \pm 0.006$ |
| **pTauS** | $0.603 \pm 0.007$ | $0.624 \pm 0.006$ | $0.621 \pm 0.005$ | $0.614 \pm 0.006$ | $0.614 \pm 0.006$ |

**Table S8.** CALVADOS parameters for phosphorylated and unphosphorylated serine and threonine (***Tesei and Lindorff-Larsen, 2023)***. pSer^HPS^ pThr^HPS^ are the previously published Perdikari parameters (***Perdikari et al., 2021)***. ^†^The fractional charge of the phosphate is calculated using experimental p*K*_*a*_ values (***Hendus-Altenburger et al., 2019)*** and the pH of the corresponding experiments, using the Henderson-Hasselbalch equation (Eq. 7). We here report charges at pH 7. ^††^Parameters for the charged serine and threonine residues used in the charge model (Ser^≈−2^ and Thr^≈−2^).

| Residue | MW [Da] | $\lambda$ | $\sigma$ [nm] | $q$ | $pK_a$ |
| --- | --- | --- | --- | --- | --- |
| **Ser** | 87.08 | 0.4625 | 0.518 | 0 |  |
| **pSer** | 165.04 | 0.0925 | 0.601 | $-1.91^\dagger$ | 6.01 |
| **Ser$^{\dagger\dagger}$** | 87.08 | 0.4625 | 0.518 | $-1.91^\dagger$ | 6.01 |
| **Asp** | 115.09 | 0.0416 | 0.558 | -1 |  |
| **Thr** | 101.11 | 0.3713 | 0.562 | 0 |  |
| **pThr** | 179.07 | 0.0013 | 0.635 | $-1.83^\dagger$ | 6.3 |
| **Thr$^{\dagger\dagger}$** | 101.11 | 0.3713 | 0.562 | $-1.83^\dagger$ | 6.3 |
| **Glu** | 129.11 | 0.0007 | 0.592 | -1 |  |
| **pSer<sup>HPS</sup>** | 165.03 | 0.162 | 0.636 | -2 |  |
| **pThr<sup>HPS</sup>** | 179.05 | 0.081 | 0.662 | -2 |  |

**Figure S1.**
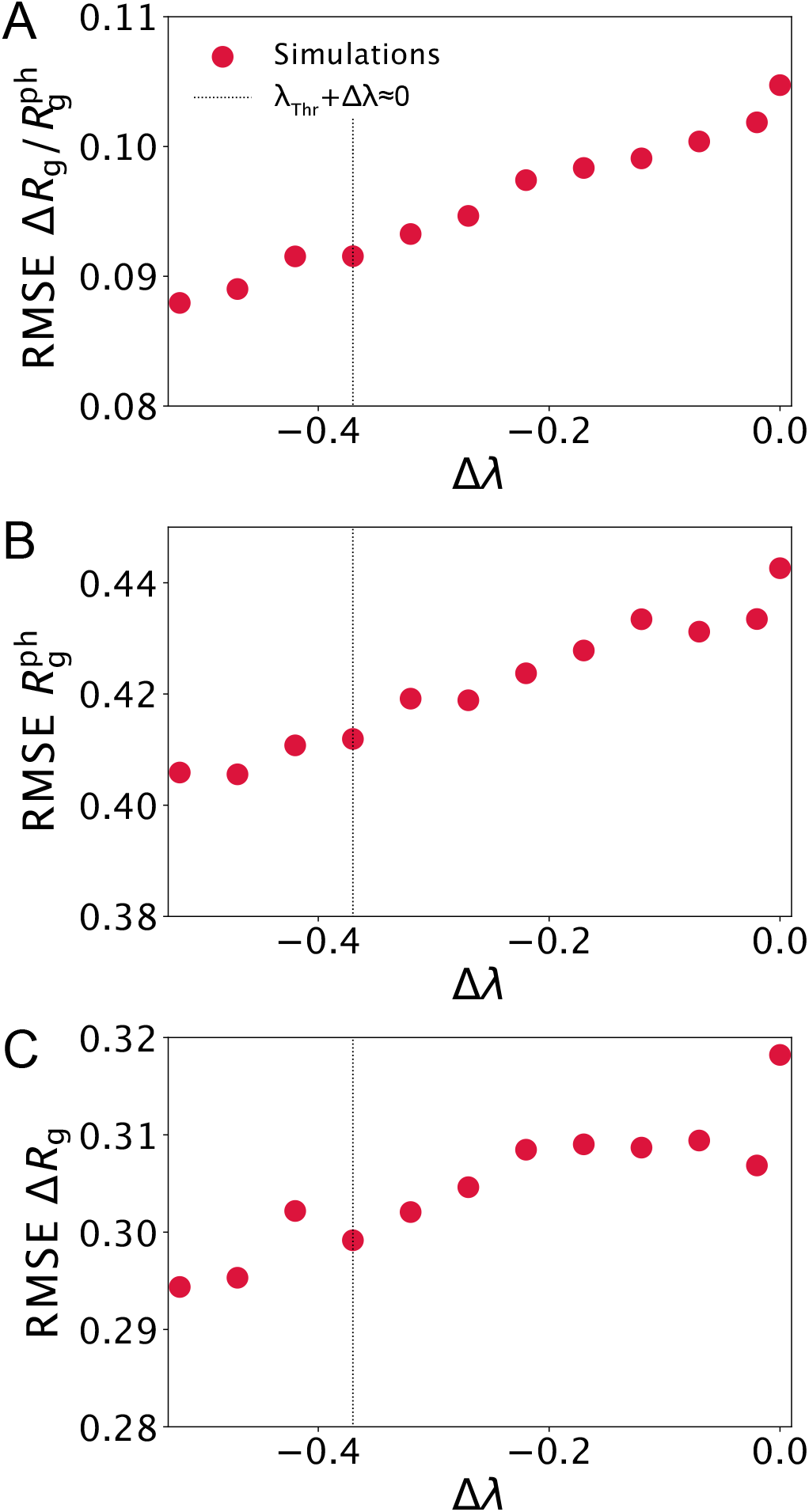
Evaluation of the phosphorylation model against experimental *R*_g_ data as a function of Δλ. (**A**) RMSE calculated for 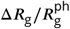. (**B**) RMSE calculated for 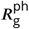. (**C**) RMSE calculated for Δ*R*_g_. Vertical dotted lines *R*g g show the Δλ resulsting in λ_pThr_ ≈ 0.

**Figure S2.**
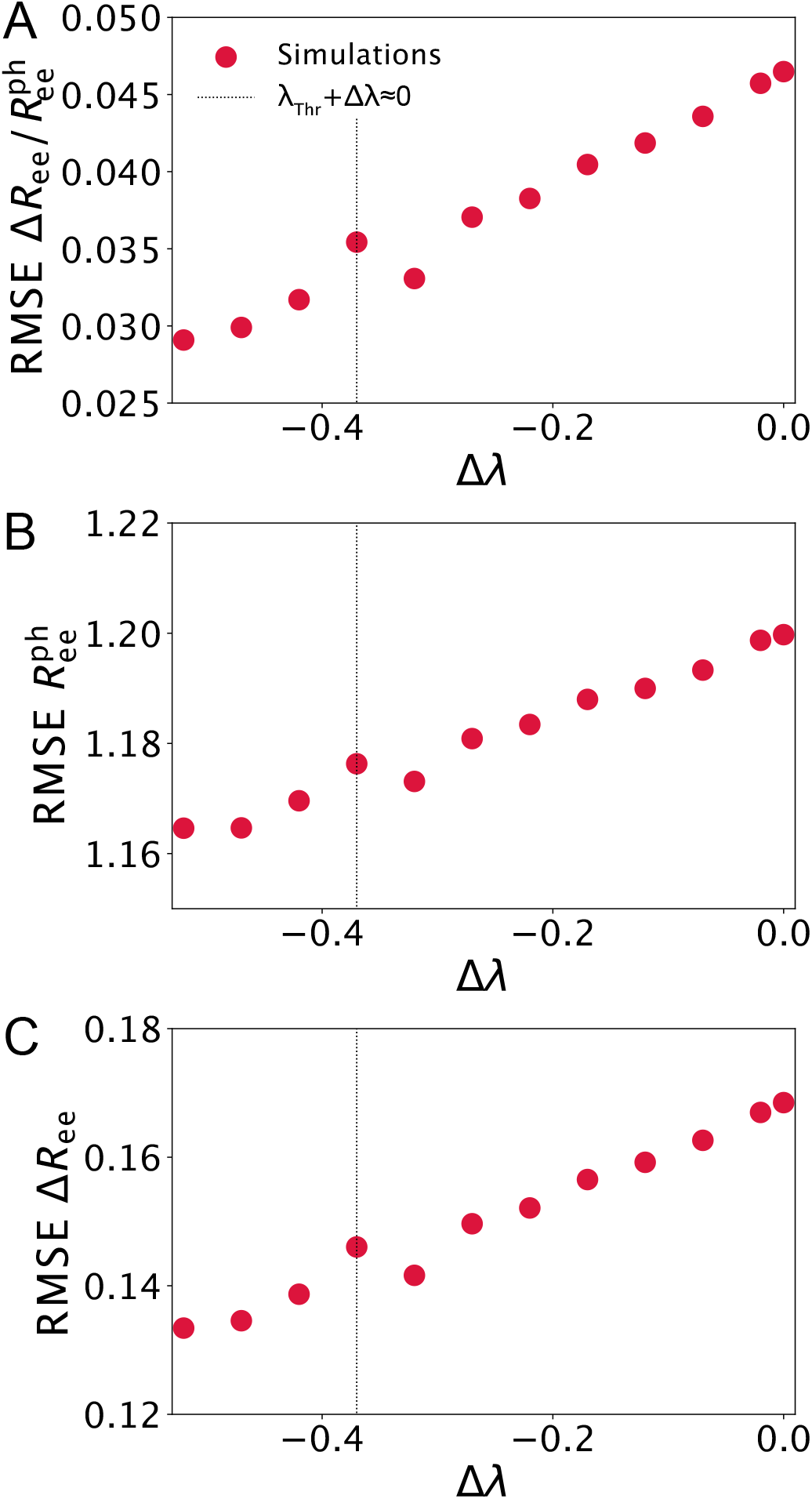
Evaluation of the phosphorylation model against experimental *R*_ee_ data as a function of Δλ. (**A**) RMSE calculated for 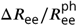. (**B**) RMSE calculated for 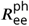. (**C**) RMSE calculated for Δ*R*_ee_. Vertical dotted *R*ee ee lines show the Δλ resulting in λ_pThr_ ≈ 0.

**Figure S3.**
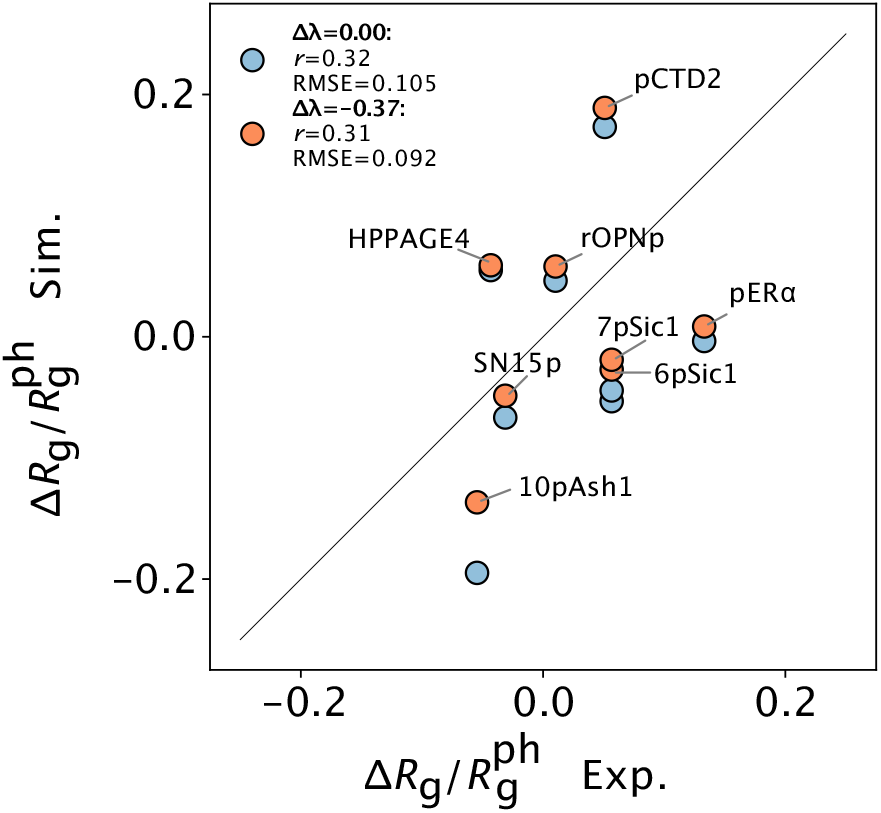
Comparison between simulated and experimental 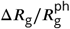 for Δλ = 0.0 (blue) and Δλ = −0.37 (orange). The solid black line shows the diagonal and the legend reports Pearson correlation coefficients, *r*, and RMSEs.

**Figure S4.**
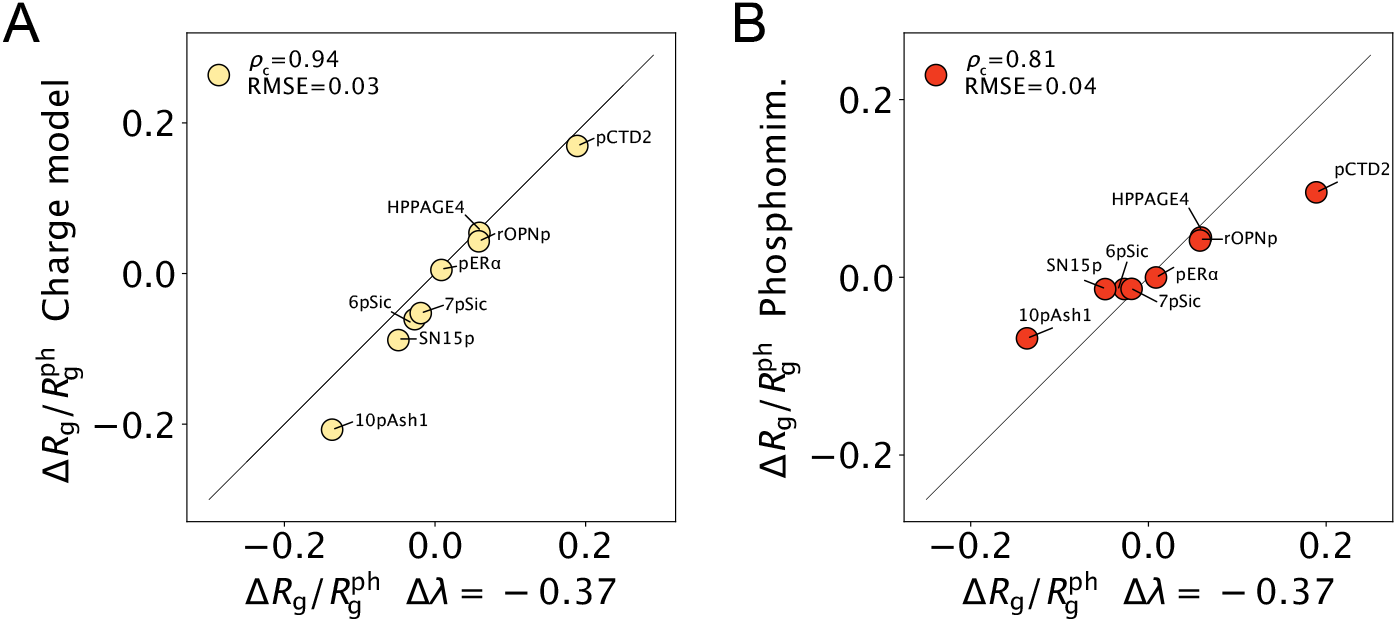
(**A**) Comparison of 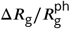 from simulations with Δλ = −0.37 and from simulations where serine and threonine residues at the phosphorylation sites are replaced with charged versions of Ser and Thr, respectively, which have a charge of ≈ −2 and the same bead sizes and stickiness as the unphosphorylated residues. (**B**) Comparison of 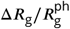 from simulations with Δλ = −0.37 and from simulations where serine and threonine residues at the phosphorylation sites are replaced with the commonly used phosphomimetics aspartate and glutamate residues, respectively. Black solid lines show the diagonals, and we report the concordance correlation coefficient, *ρ*_c_, (***Lin, 1989)*** and the RMSE in the legends.

## Notes

https://github.com/KULL-Centre/_2025_rauh_phosphorylation

### Competing Interest Statement

KL-L holds stock options in and is a consultant for Peptone Ltd.

